# Characterization of Social and Repetitive Behaviors of Mllt11/Af1q/TcF7c Conditional Knockout Mice

**DOI:** 10.1101/2024.07.31.606037

**Authors:** Emily A. Witt, Danielle Stanton-Turcotte, Pristine Garay, Jinrong Ge, Angelo Iulianella

## Abstract

*Mllt11* (myeloid/lymphoid or mixed-lineage leukemia translocated to chromosome 11; also known as *Af1q/TcF7c*) has been identified as a novel regulator of neural development, playing a role in the migration and outgrowth of cortical projection neurons. We previously reported that the conditional inactivation of the *Mllt11* gene in the mouse superficial cortex resulted in reduced connectivity of the corpus callosum and white matter fiber tracts, resulting in reduced cortical thickness. However, the behavioral consequences of *Mllt11* loss are unknown. Callosal abnormalities are thought to be present in 3-5% of all neurodevelopmental disorders and reduced corpus callosum volume correlates with core symptoms of autism spectrum disorder (ASD) in humans. Cortical thickness dysregulation is likewise shared among various neurodevelopmental disorders including ASD. We therefore investigated the behavioral consequences of conditional knockout of *Mllt11* in upper cortical layer 2/3 projection neurons using transgenic *Cux2^iresCre^* mice. Utilizing tasks designed to reflect core ASD symptoms, we examined the behaviors of both male and female conditional knockout animals. These tests included olfaction habituation/dishabituation, three-chambered social approach, marble burying, and nestlet shredding. We found sex-dependent disruptions in social preference and nestlet shredding in animals lacking *Mllt11*, with the female mice presenting with more disruptions than the males. Understanding the behavioral phenotype associated with genes of interest, specifically in the context of sex differences, is crucial to individualized treatment for neurodevelopmental disorders.

## INTRODUCTION

The genesis of the mammalian brain is dependent upon the regulation of cytoskeletal proteins that promote the migration of neurons, extension of axonal growth cones and dendrites, and ultimately functional, synaptically-coupled networks^1,2^. Specifically, axons and dendrites are underlain by a structural network of microtubules (MTs), actin, intermediate filaments, and associating proteins that change dynamically to promote neurite outgrowth and synaptogenesis^1^. It is therefore not surprising that mutations that alter MTs and associated proteins are implicated in neurodevelopmental disorders (NDDs). Our lab has been investigating a neuronally-restricted cytoskeletal-associated protein called Mllt11 (myeloid/lymphoid or mixed-lineage leukemia translocated to chromosome 11; also known as Af1q, or Tcf7c). A previous report identified a novel copy number variant (CNV) in *Mllt11* in an individual with ASD. The affected chromosomal region included a deletion of MLLT11 among other genes, although at the time, as the authors mentioned “the function..[of MLLT11]..is either unrelated to brain function or poorly characterized”^3^. Our investigations have since provided evidence for the role of Mllt11 in neural development, regulating both migration of cortical neurons, and neuritogenesis, which is required for the establishment of neuronal connectivity.

We found that *Mllt11* is enriched in developing neurons during the formation of the upper layers (UL) 2/3. Thus, we generated a conditional knockout (cKO) mouse that inactivated *Mllt11* in the UL 2/3 of the cerebral cortex using the *Cux2^iresCre^* driver mouse line^4–6,7^. We found that *Cux2^iresCre^*; *Mllt11^flox/flox^* cKO animals displayed a thinning of the cortex, reduced complexity of superficial cortical projection neuron (CPN) arborization patterns and reduced axonal projections across the corpus callosum at embryonic timepoints^8^; consistent with anatomical abnormalities reported in various NDDs including ASD and schizophrenia (SZ)^9–15^. Although divided into distinct disorders, NDDs converge on various levels in symptoms, behavior, biomarkers, and genetic risk^16^ and are often co-diagnosed^17–20^. Two such convergences are the development of the corpus callosum and altered cortical thickness^21^. Two hundred million callosal axons integrate and transfer information between the two hemispheres in the human brain^22,23^. This communication is integral for the various functions of the cerebral cortex including management of social and emotional stimuli; as such, abnormalities in the corpus callosum are thought to be present in 3-5% of all neurodevelopmental disorders^24,25^. Differences in cortical thickness, likewise, are shared among at least 6 common NDDs^21,26,27^. *Mllt11* has also been found to be dysregulated in fetal mice following maternal valproic acid (VPA) administration - a common mouse model used to study ASD^28,29^. Given that *Mllt11* cKO mouse brains share histological abnormalities with individuals with NDDs, its dysregulation following VPA, and the CNV of the region containing *Mllt11* in an ASD patient, we hypothesized that Mllt11 loss of function in our *Cux2^iresCr^*^e^-driven *Mllt11* cKO mice would display ASD behavioral symptomology. To investigate this, we selected behavioral tasks which reflect the core human ASD symptomology^30^: communication via the olfactory habituation dishabituation task; sociability via the three-chamber social preference task; and examination of repetitive/compulsive behaviors via marble burying, nestlet shredding and scoring of behaviors such as digging and grooming during other tasks. Additionally, we confirmed that the presentation of altered thickness of the cortex and the corpus callosum was present postnatally.

## MATERIALS AND METHODS

### Animals

All experiments described herein were performed according to approved IACUC protocols at Dalhousie University. Mice were generated as previously described^8^. Briefly, *Mllt1^flox/flox^*; *Rosa26^TdTomato^ mice* were crossed with *Mllt11^flox/+^;Cux2^iresCre^;Rosa26^TdTomato^*mice to generate the following mice for testing: littermate Control animals – *Mllt11^flox/+;^Rosa26^TdTomato^,* Heterozygous animals – *Mllt11^flox/+^; Cux2^iresCre^;Rosa26^TdTomato,^*and finally conditional knockouts (cKO) – *Mllt11^flox/flox^;Cux2^IRESCre^;Rosa26^TdTomato^.* As these crosses did not allow for littermate controls that express Cre alone, Cre control animals were generated by crossing *Rosa26^TdTomato+/−^* mice with *Cux2^iresCre^;Rosa26^TdTomato^*^+/−^ mice. These crosses generated the following mice for testing: WT - *Mllt11^+/+^;Rosa26^TdTomato^*and Cre – *Mllt11^+/+^;Cux2^iresCre^;Rosa26^TdTomato^*. Both male and female mice of the correct genotype were used for each experiment. Animals were housed with sex matched littermates. Animals were ear clipped on P8 and ear clips were used for genotyping. The offspring produced were all genotyped via PCR as described previously^8^.

### Behavioral Analysis

All behavioral experiments took place during the light cycle between (7am-7pm). A full timeline of the experiments performed can be seen in **Figure. 1**. Animals were weaned on postnatal day (P) 23. One week prior to initiation of testing, animals were brought daily to the testing room for habituation to the room and handling. First, the animals were left for 10 minutes to habituate to the testing room. After 10 minutes, the animals were gently handled, weighed, and then returned to their home cage in the animal colony room.

**Figure 1:**
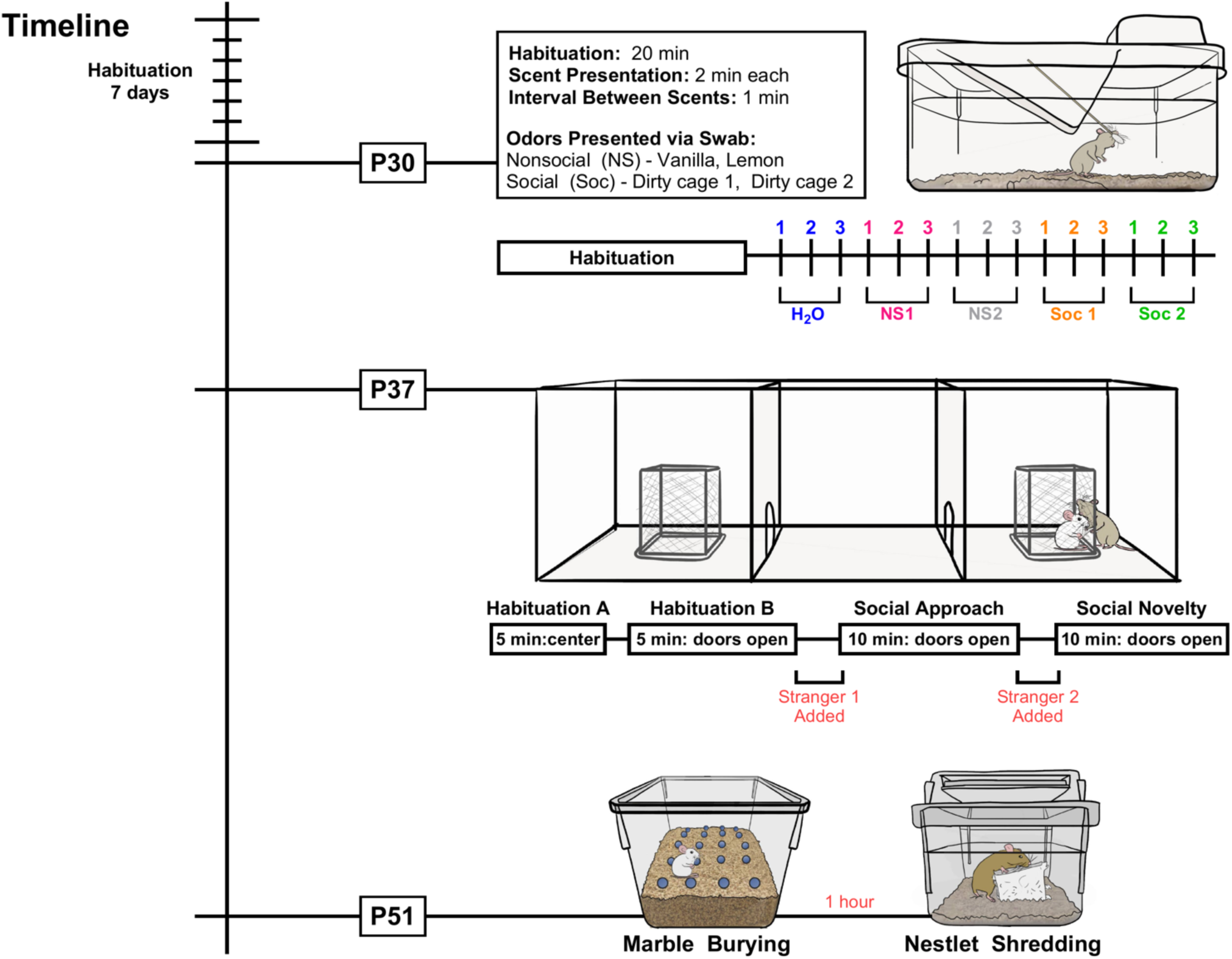
Timeline of Experiments. Timeline of behavioral experiments; described in detail within the methods section.

### Olfaction Habituation/Dishabituation

This test was performed on postnatal day 30 (P30) and utilized the procedures outlined by Yang and Crawley ^31^. Animals were placed into a clean, empty cage with corn cob bedding and brought to the testing room. A 20-minute period of habituation was initiated by insertion of a clean, dry cotton swab into the cage. Following habituation, sequences of three swabs dipped in identical odors were inserted into the cage for 2 minutes each with 1-minute intertrial intervals. The odors are as follows: (1) distilled water, (2) vanilla extract 1:100, (3) lemon extract 1:100, (4) swabs soaked in distilled water and run in a zig-zag pattern along the bottom of a three-day old unfamiliar, dirty mouse cage of each sex acquired fresh the day of testing. Presentation of the odors was done in the following order: three distilled water, three non-social odor 1, three non-social odor 2, three social odor 1, and three social odor 2, and the order of specific odors within this paradigm was randomized and balanced across sexes and genotypes. Every subject was exposed to novel social odor from two different dirty cage sources. An experimenter blinded to the genotypes scored time spent sniffing the swab with a stopwatch. Video recordings were acquired and scored for repetitive behaviors (see below).

### Three-Chamber Social Approach

This test was performed on postnatal day 37 (P37). Our protocol was based upon a protocol from Kaidanovich-Beilin, Lipina, Vukobradovic, Roder and Woodgett ^32^. Briefly, animals were first placed in a clean empty cage with corncob bedding and brought to the testing room. The mouse being tested was then placed in the center of the three-chambered apparatus with the two openings closed for 5 minutes (Habituation A). After 5 minutes, the doors to each side of the chamber were opened and the mouse was able to freely roam the entire arena for an additional 5 minutes (Habituation B). The right and left side of the chamber contained empty, upside-down wire mesh pencil cups in the center. After 5 minutes, the doors to each peripheral chamber were closed when the testing mouse returned to the center chamber. A novel age- and sex-matched mouse was then placed inside the pencil cup in one of the adjacent chambers. (All novel mice were habituated to the enclosures the day before testing. Briefly described, the novel mice were placed in the pencil cup for 10 minutes each, then returned to the home cage in the testing room for 10 minutes, then returned to the pencil cup for 10 minutes. This was done until each novel mouse completed three 10-minute cycles in the enclosure.) Both doors of the chamber were then opened, and the testing mouse was able to freely explore the apparatus for 10 minutes. At the end of 10 minutes, the doors to each chamber were closed once the mouse returned to the center chamber. A second novel mouse was then placed inside the previously empty pencil cup. The doors were then removed, and the mouse was able to freely roam the entire arena for an additional 10 minutes. Video recordings were acquired, and time spent sniffing the pencil cups, and time spent in each area of the apparatus were recorded for each segment of the experiment (Habituation A: 5 min, Habituation B: 5 min, Stranger 1: 10 min, Stranger 2: 10 min) by an experimenter blind to the mouse genotype.

### Marble Burying and Nestlet Shredding

This test was performed on postnatal day 51 (P51). Our protocol was based upon a previously published protocol from Angoa-Perez, Kane, Briggs, Francescutti and Kuhn ^33^. The mice were brought to the testing room as described previously and placed into corncob lined clean cages with a single pre-weighed cotton nestlet square in the center of the cage. Mice were left for 30 minutes and the amount of unshredded nestlet was weighed. The mice were then returned to their home cage in the colony room for 1 hour. During this time, standard rat cages were filled with 5 cm of fresh corn cob bedding. Glass marbles were placed in a grid pattern with 5 rows of 4 marbles evenly spaced. After 1 hour, mice were returned to the testing room and placed in the prepared rat cage and allowed to freely roam and interact with the marbles for 30 minutes. The number of buried marbles was recorded by an experimenter blind to the genotypes at the end of 30 minutes.

### Video scoring & Behavioral Definitions

Video scoring was done by experimenters blinded to the genotypes of the mice. Each scorer was responsible for measuring a particular behavior to avoid effects of interrater reliability. In the olfactory habituation/dishabituation task, investigation was defined as when the animal was oriented towards the cotton swab with the nose of the animal being 2cm or closer to the cotton tip. Investigation of the wooden portion of the stick was not counted. Investigation was defined similarly in the three-chamber task, as when the animal was oriented towards the pencil cup with the animal’s nose being 2cm or closer to the pencil cup. Grooming in all tests was defined as time of active self-cleaning that utilizes friction movements with the paws directed at the body or tail or self-cleaning utilizing the mouth on the body or tail. Digging was defined as time when the animal was actively utilizing their paws or head to move corncob bedding. Rearing in all tasks was defined as the lifting of the forepaws from the ground and standing in an upright position on the hindlimbs. Freezing was defined as total immobility except for respiratory movement. For the three-chamber task, total freezing was recorded as being the sum of the freezing time measured in each individual section of the experiment (Freezing during Habituation A + freezing during Habituation B + freezing during Stranger 1 + freezing during Stranger 2 = Total Freezing).

### Histology and Immunohistochemistry

For histological analysis, a separate cohort of mice not subjected to behavioral experimentation underwent transcardiac perfusions with PBS followed by 4% PFA on postnatal day 30 (P30). Brains were removed and fixed in 4% PFA overnight at 4°C. The following day, the brains were equilibrated progressively in 15% and 30% sucrose in PBS for cryoprotection. 20µm sections were collected serially via cryosectioning and were immunostained with DAPI and mouse anti-Neurofilament 2H3 (1:200; DSHB, University of Iowa). Anti-mouse Alexa Fluor 647-conjugated IgG (1:1500; Invitrogen) secondary was used to detect the primary.

### Microscopy

Sections were imaged on a Zeiss Axio Observer Fluorescence microscope and a Hamamatsu Orca Flash v4.0 digital camera at 10x magnification. Images were processed with Zeiss Zen Software. Per individual, three images of the corpus callosum as well as three images of the cortex were acquired. Measurements of the corpus callosum, and cortex were collected using ImageJ (FIJI).

### Statistical Analysis

Analysis was performed using Prism 10 software. Two-way ANOVAs were performed when comparing male or female *Mllt11^flox/+^*, *Mllt11^flox/+^;Cre*, and cKO animals on the cumulative olfaction habituation/dishabituation task and the three-chamber social task. One-way ANOVAs were performed when comparing male or female *Mllt11^flox/+^*, *Mllt11^flox/+^;Cre*, and cKO animals marble burying, nestlet shredding, grooming, digging, rearing, total freezing, corpus callosum width and cortex depth. Homogeneity of variance was confirmed using the Brown-Forsythe Test. For Cre control cohorts, two-way ANOVAs were performed when comparing male or female WT vs Cre animals on the olfaction habituation/dishabituation task and the three-chamber social task. Unpaired T-Tests were performed when comparing male or female WT vs Cre animals marble burying, nestlet shredding, grooming, digging, rearing, total freezing, corpus callosum width, and cortex depth. For multiple comparisons, Sidak’s test was used to explore significant effects and Tukey’s test was used to explore interactions found between two or more genotypes when the number of comparisons per column family was greater than one and Uncorrected Fisher’s LSD was used to explore interactions found between two genotypes when the number of comparisons per column family was one. For all graphs, error bars reflect the standard error of the mean (SEM) and ns= not significant; * = p<0.05; ** = p <0.01; *** = p<0.001; **** = p<0.0001).

## RESULTS

### Female cKO animals display reduced time spent with social odors

Olfactory ability and interest in social odors were assessed by evaluating habituation/dishabituation and time spent investigating social and non-social odors among the various genotypes. All three genotypes (*Mllt11^flox/+^*, *Mllt11^flox/+^;Cre*, and cKO) in males and females displayed habituation to each of the tested odors as consecutive presentations of odors resulted in reduced investigation **(Fig. 2a-b).** Furthermore, all three genotypes dishabituated as there was an increase in investigation time with each presentation of a novel odor **(Fig. 2a-b)**. We then evaluated the interest in social odors by examining the cumulative time spent investigating the social odors in each genotype **(Fig. 2c-d)**. Two-way ANOVA revealed no significant interaction of genotype and odor between the three male genotypes with the time spent with social or non-social odors (F(2,31)=0.002992; p=0.9970; **Fig. 2c**). There was a significant effect of odor (F(1,31)=50.89; p <0.0001) and Sidak’s multiple comparisons revealed all three genotypes spent significantly more time investigating the social odors (*Mllt11^flox/+^* p=0.0005; *Mllt11^flox/+^;Cre*: p=0.0018; cKO p=0.0005; **Fig 2c**). However, two-way ANOVA within the females revealed a significant interaction of genotype and odor (F(2,34)= 3.881; p=0.0303) and Tukey’s multiple comparisons found that the female cKO animals spent significantly less time investigating the social odors than the *Mllt11^flox/+^* (p=0.0474) and Mllt11^flox/+^;Cre (p=0.0007) animals **(Fig. 2d)**. As with the male mice, all three female genotypes spent significantly more time investigating the social odors than the nonsocial odors (*Mllt11^flox/+^* p<0.0001; *Mllt11^flox/+^;Cre*: p<0.0001; cKO p =0.0342; **Fig. 2d**).

**Figure 2:**
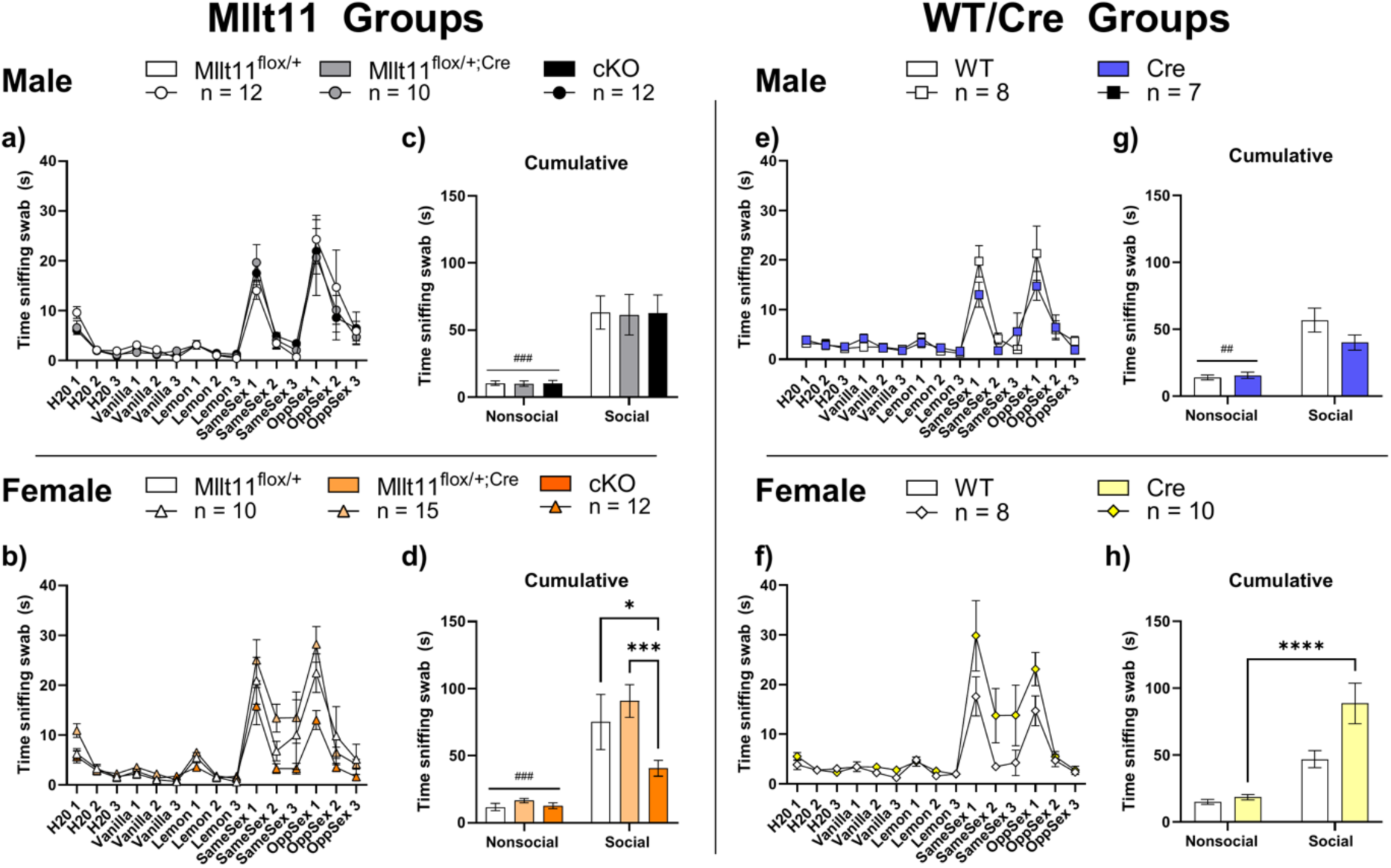
Olfaction Habituation/Dishabituation. Graphical representation of the recorded time spent sniffing the swab during the olfactory habituation/dishabituation test. **a)** Time spent sniffing swabs in the male *Mllt11^flox/+^, Mllt11^flox/+^;Cre*, and cKO genotype. **b)** Time spent sniffing swabs in the female *Mllt11^flox/+^, Mllt11^flox/+^;Cre,* and cKO genotypes. **c)** Cumulative time spent sniffing the two nonsocial odors vs the two social odors was compared in the male *Mllt11^flox/+^, Mllt11^flox/+^;Cre*, and cKO genotypes. **d)** Cumulative time spent sniffing the two nonsocial odors vs the two social odors was compared in the female *Mllt11^flox/+^, Mllt11^flox/+^;Cre*, and cKO genotypes. **e)** Time spent sniffing swabs in the male WT and Cre genotypes. Time spent sniffing swabs in the female WT and Cre genotypes. **g)** Cumulative time spent sniffing the two nonsocial odors vs the two social odors was compared in the male WT and Cre genotypes. **h)** Cumulative time spent sniffing the two nonsocial odors vs the two social odors was compared in the male WT and Cre genotypes. (For all graphs, ### = All three genotypes cumulative non-social time was significantly less than cumulative social time (p<0.05). ## = All two genotypes cumulative non-social time was significantly less than cumulative social time (p<0.05); * = p<0.05; *** = p<0.001; **** = p<0.0001).

To ensure that the effects in the *Mllt11^flox/+^;Cre* and cKO animals were not due to Cre expression, we performed the same tests in our Cre control genotypes **(Fig 2e-h)**. Beginning with males, two-way ANOVA did not reveal a significant interaction of genotype and odor (F(1,13)=3.282; p=0.0932; **Fig. 2g**) but did reveal an effect of odor (F(1,13)=45.37; p<0.0001) on investigation time. Sidak’s multiple comparisons revealed that both genotypes spent significantly more cumulative time investigating the social odors (WT p<0.0001; Cre p=0.0100; **Fig. 2g**, significance indicated by ##). Similarly, between the females, a two-way ANOVA did not reveal a significant interaction of genotype and odor (F(1,16)=4.478; p=0.0504) on investigation time **(Fig. 2h)**. There was an effect of odor (F(1,16)=31.64; p<0.0001) on investigation time. Sidak’s multiple comparisons revealed that although both genotypes investigated the social odor more, the Cre genotype but not the WT genotype spent significantly more cumulative time with the social odors (WT p=0.0624; Cre p<0.0001; **Fig. 2h**). This suggests that cKO of *Mllt11* in females but not males resulted in reduced interest in social odors and that this difference is due to loss of both *Mllt11* copies and not simply expression of Cre as the effects of Cre expression alone were not identical to those of the *Mllt11* cKO genotypes.

### Sex-specific disruptions in social preference, novelty, and freezing

We next evaluated the social preference of the mice using the three-chamber social task. Beginning with the males, two-way ANOVA revealed a significant interaction between genotype and chamber side (F(2,32)=3.868; p=0.0313; **Fig. 3a**) on time spent in the chamber. Sidak’s multiple comparisons showed that the *Mllt11^flox/+^;Cre* males and the cKO males both did not display a significant preference for the chamber containing the stranger mouse (*Mllt11^flox/+^;Cre* p =0.5972; cKO p=0.1499; **Fig. 3a**). The *Mllt11^flox/+^* males however did display a significant preference for the chamber containing the stranger mouse (p=0.0001; **Fig. 3a**). As the previous test found no differences in investigation of social odors between the males, we next evaluated the investigation behavior of the males (defined as sniffing the pencil cup) (**Fig. 3b)**. In alignment with the social odor presentation test, two-way ANOVA revealed no significant interaction between genotype and chamber side (F(2,32)=2.785; p=0.0768) on time spent, but did reveal a significant effect of chamber side (F(1,32)= 8.68; p<0.0001; **Fig. 3b**) on time spent. Sidak’s multiple comparisons revealed that all three genotypes spent significantly more time investigating the pencil cup containing the stranger than the empty pencil cup (*Mllt11^flox/+^* p<0.0001; *Mllt11^flox/+^;Cre*: p=0.0023; cKO p=0.0002; **Fig. 3b**). Demonstrating that all three genotypes distinguished and attended to the presence of the stranger mouse **(Fig. 3b)**.

**Figure 3:**
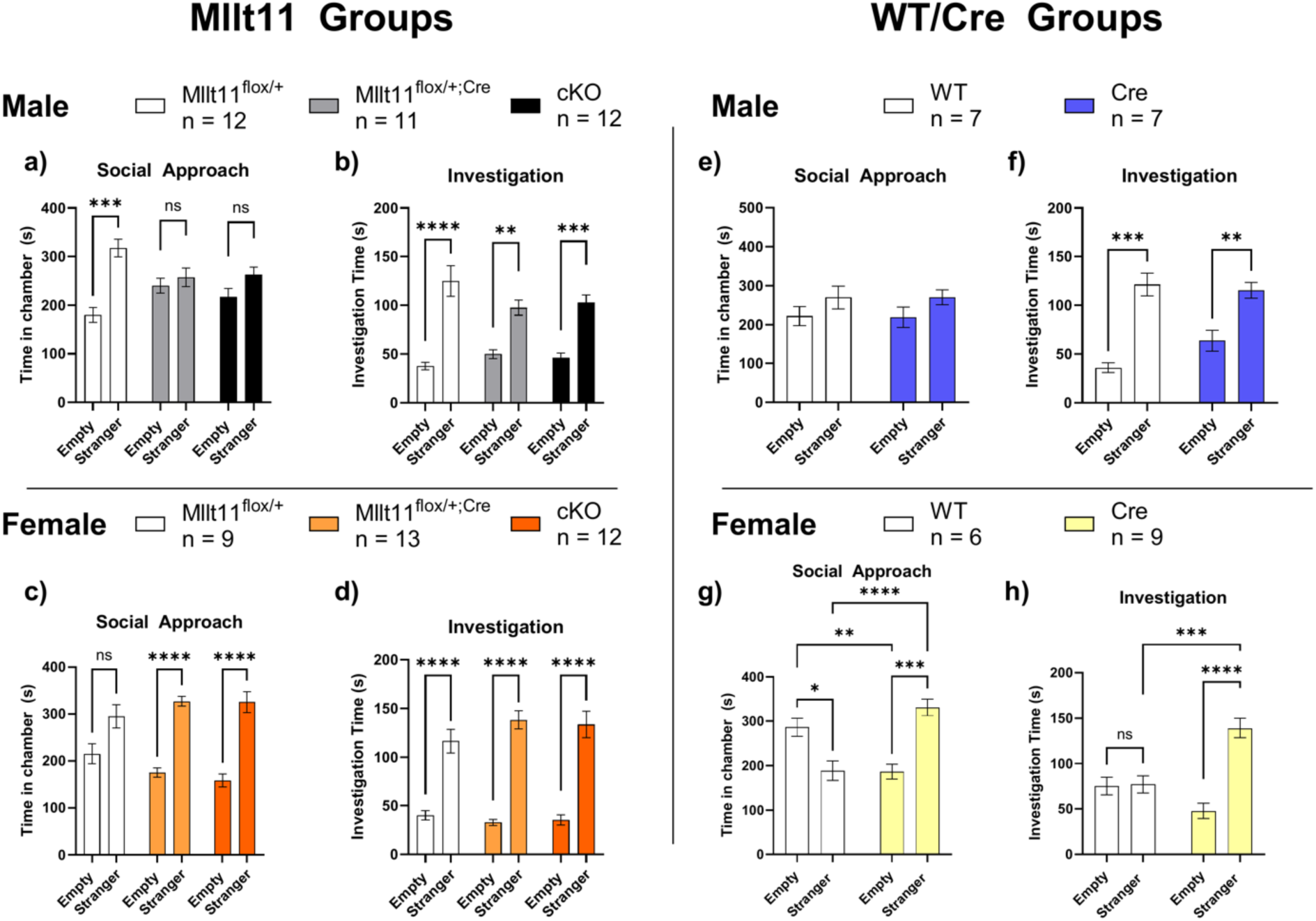
Three-Chamber Social Approach. Graphical representation of the data recorded during the Social Approach portion of the three-chamber task. This is the stage in which one side of the chamber remains empty while the other side contains the first stranger mouse. The time spent by the mice in either the ‘Empty’ chamber which contained just an empty pencil cup or in the ‘Stranger’ chamber which contained the stranger mouse within a pencil cup was recorded and compared between the **a)** male *Mllt11^flox/+^, Mllt11^flox/+^;Cre*, and cKO genotypes, **c)** female *Mllt11^flox/+^, Mllt11^flox/+^;Cre*, and cKO genotypes, **e)** male WT and Cre genotypes, female WT and Cre genotypes. As a measure of investigation, the time spent by the mice actively sniffing either the ‘Empty pencil cup or the pencil cup containing the ‘Stranger’ was recorded and compared between the **b)** male *Mllt11^flox/+^, Mllt11^flox/+^;Cre*, and cKO genotypes, **d)** female *Mllt11^flox/+^, Mllt11^flox/+^;Cre*, and cKO genotypes, **f)** male WT and Cre genotypes, **h)** female WT and Cre genotypes. (For all graphs ns = not significant; * = p<0.05; ** = p<0.01; *** = p<0.001; **** = p<0.0001).

Among the females, two-way ANOVA found no interaction between genotype and chamber side (F(2,31)=1.951; p=0.1591) on time spent in chamber, but a significant effect within genotype in the time spent in either peripheral chamber (F(1,31)=53.64; p<0.0001; **Fig. 3c**). Sidak’s multiple comparisons test showed that the *Mllt11^flox/+^;Cre* and cKO mice all displayed a significant preference for the chamber containing the stranger (*Mllt11^flox/+^;Cre* p<0.0001; cKO p<0.0001) while the *Mllt11^flox/+^* did not (p=0.0825; **Fig. 3c**). Evaluation of investigation behavior in the females revealed no significant interaction between genotype and chamber side (F(2,31)=1.253; p=0.2997) on time spent, but did reveal a significant effect of chamber side within genotype (F(1,31)=156.4; p<0.0001; **Fig. 3d**) on time spent. Sidak’s multiple comparisons revealed similarly to the males that all three female genotypes spent significantly more time investigating the pencil cup containing the stranger (Mllt11^flox/+^ p<0.0001; *Mllt11^flox/+^;Cre*: p<0.0001; cKO p<0.0001) once again demonstrating that, while they did not display social preference, they did display preferential investigation of the presence of the stranger mouse **(Fig. 3d)**.

To exclude any effects of Cre, we performed the same experimentation on cohorts of Cre controls. Neither male genotype displayed a significant preference for the chamber containing the stranger (F(1,12)=0.002267; p=0.9628; **Fig. 3e**). As this lack of preference mirrors that of the male *Mllt11^flox/+^;Cre* males and the cKO, we believe the lack of preference cannot be explained by the lack of one or both copies of *Mllt11* alone. Examination of investigation behavior in the males revealed a significant effect of chamber side within genotype (F(1,12)=42.56; p<0.0001) on time spent, but no interaction between genotype (F(1,12)=2.599; p=0.1329). Sidak’s multiple comparisons showed that both the WT and the Cre animals spent more time investigating the pencil cup containing the stranger (WT p=0.0002; Cre p=0.0092; **Fig. 3f**). Within the female Cre control cohort, there was a significant interaction between genotype and chamber side (F(1,13)=20.38; p=0.0006) on time spent. Uncorrected Fisher’s LSD multiple comparisons revealed that the WT animals displayed a significant preference for the empty chamber (p=0.0351) while the Cre females displayed a significant preference for the chamber containing the stranger (p=0.0009; **Fig. 3g)**. We then examined the amount of time the WT/Cre females spent investigating each pencil cup. We found that the Cre genotype, but not the WT genotype, spent significantly more time investigating the pencil cup containing the stranger (p<0.0001) and that the Cre animals spent more time than the WT animals investigating the pencil cup containing the stranger (p=0.0002; **Fig. 3h**). As these results are not what we saw in our three female Mllt11 genotypes, the effects seen in our Mllt11 genotypes can be attributed to *Mllt11* loss.

Examination of the novelty stage of the task identified some interesting results. In the male Mllt11 experimental genotypes, two-way ANOVA found no significant interaction between genotype and chamber side (F(2,32)=0.7323; p=0.4887) on time spent, but did find a significant effect of chamber side within genotypes (F(1,32)=12.95; p=0.0011) on time spent. Sidak’s multiple comparisons showed that only the *Mllt11^flox/+^;Cre* males spent significantly more time in the novel stranger side (p=0.0301; **Fig. 4a**). However, examination of investigation times showed that, while there was no significant interaction between genotype and pencil cup chamber side (F(2,32)=1.695; p=0.1997) on time spent investigating the associated pencil cup, there was a significant effect of pencil cup chamber side within genotype (F(1,32)=44.49; p<0.0001) on time spent investigating the associated pencil cup that indicated via Sidak’s multiple comparisons that only the *Mllt11^flox/+^;Cre* and cKO spent significantly more time investigating the pencil cup containing the novel stranger (*Mllt11^flox/+^;Cre* p=0.0002; cKO p=0.0002; **Fig. 4b**).

**Figure 4:**
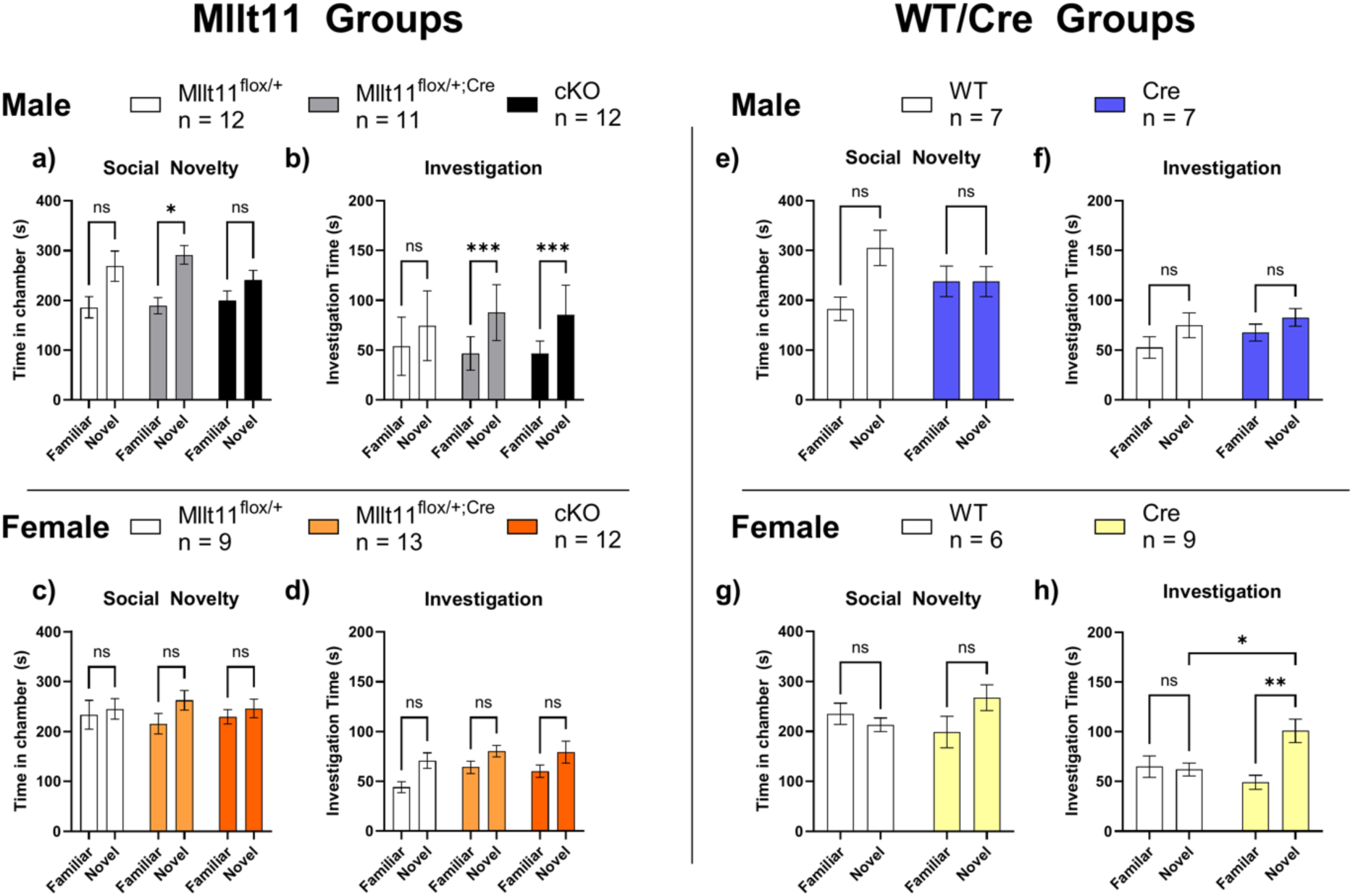
Three-Chamber Social Novelty. Graphical representation of the data recorded during the Social Novelty portion of the three-chamber task. This is the stage in which a second Stranger mouse – the ‘novel’ mouse is added to the pencil cup in the previously empty side of the chamber. The time spent by the mice in either the ‘Familiar’ chamber, which contained the same ‘Stranger’ mouse from the Social Approach portion contained in in a pencil cup, or in the ‘Novel’ chamber which contained the ‘Novel’ mouse within a pencil cup, was recorded and compared between the **a)** male *Mllt11^flox/+^, Mllt11^flox/+^;Cre*, and cKO genotypes, **c)** female *Mllt11^flox/+^, Mllt11^flox/+^;Cre*, and cKO genotypes, **e)** male WT and Cre genotypes, **g)** female WT and Cre genotypes. As a measure of investigation, the time spent by the mice actively sniffing either the pencil cup containing the ‘Familiar’ mouse or the pencil cup containing the ‘Novel’ mouse was recorded and compared between the **b)** male *Mllt11^flox/+^, Mllt11^flox/+^;Cre*, and cKO genotypes, **d)** female *Mllt11^flox/+^, Mllt11^flox/+^;Cre,* and cKO genotypes, **f)** male WT and Cre genotypes, **h)** female WT and Cre genotypes. (For all graphs ns= not significant * = p<0.05; ** = p<0.01; *** = p<0.001; **** = p<0.0001).

In the females, there was not a significant interaction between genotype and chamber side (F(2,31)=0.2531; p=0.7780) on time spent and no significant effect within genotype and chamber side (F(1,31)=1.227; p=0.2765) on time spent. Examination of investigation time found no significant interaction between genotype and pencil cup chamber side (F(2,31)=0.2749; p=0.7615) on time spent investigating the associated pencil cup, but did find an effect within genotype (F(1,31)=12.94; p=0.0011) on time spent investigating the associated pencil cup, however Sidak’s multiple comparisons revealed that there were no differences in time spent investigating the novel vs familiar mouse in any of the three Mllt11 experimental genotypes (*Mllt11^flox/+^*p=0.0631; *Mllt11^flox/+^;Cre* p=0.2393; cKO p=0.1553; **Fig. 4d**).

We performed the same tests in our male and female WT/Cre animals to determine if the effects we saw were due specifically to the loss of *Mllt11* as opposed to Cre expression in cortical projection neurons. First, WT/Cre males showed no significant effect between (F(1,12)=2.171; p=0.1663) or within (F(1,12)=2.126; p=0.1705) genotype with regard to time spent in chamber sides **(Fig. 4e)**. Similarly, there were no effects between (F(1,12)=2.171; p=0.1143) or within (F(1,12)=3.182; p=0.0997) genotypes with regard to investigation times **(Fig. 4f)**. As the effects seen in the Mllt11 experimental genotypes were not replicated in the WT/Cre control genotypes, this indicates that the effects in the Mllt11 males are due to loss of *Mllt11* and not Cre expression. Examination of our WT/Cre females likewise found no significant effects between (F(1,13)=1.506; p=0.2415) or within (F(1,13)=0.4029; p=0.5366) genotypes and time spent on either chamber side (**Fig. 4g)**. When examining the investigation times, there was a significant interaction between genotype and pencil cup chamber side (F(1,13)=6.327; p=0.0258) and within genotypes (F(1,13)=5.085; p=0.0420; **Fig. 4h**) on time spent investigating the associated pencil cup. Uncorrected Fisher’s LSD multiple comparisons found that the WT females did not spend a significantly different amount of time investigating the novel mouse vs the familiar (p=0.8692) but the Cre females did spend significantly more time investigating the novel mouse than the familiar (p=0.0023; **Fig. 4h**). Furthermore, the investigation time towards the novel mouse was found to be significantly higher in the Cre genotype than the WT genotype (p=0.2727; **Fig. 4h**). These effects were absent in the *Mllt11^flox/+^, Mllt11^flox/+^;Cre,* and cKO cohorts indicating that the results in the experimental genotypes were due to Cre expression alone.

We then compared freezing times between genotypes. One-way ANOVA revealed no significant difference between males’ average number of bouts (F(2,32)=1.711; p=0.1969; **Fig. 5a**) average length of freezing bout (F(2,32)=2.027; p =0.1483; **Fig. 5b**) across the entire session. To visualize whether the amount of freezing would correlate in any way to the addition of a stranger mouse, we quantified freezing specifically within each portion of the three-chamber task performed on P37 (Habituation A, Habituation B, Approach, Novelty). Two-way ANOVA did not find a significant interaction of genotype and portion of the three-chamber task (F(6,96)=1.223; p=0.3011; **Fig. 5c**) but did find an effect within genotypes (F(3, 96); p=0.0164) on time spent freezing. Tukey’s multiple comparisons did not reveal any differences **(Fig. 5c)**. Within the females, there was not a significant difference in the average number of freezing bouts (F(2,31)=1.474; p=0.2445; **Fig. 5d**) however, there was a significant difference within the length of freezing bouts (F(2,31)=4515; p=0.0190) with multiple comparisons revealing that the cKO females average length of each freezing bout was significantly longer than the *Mllt11^flox/+^* (p=0.0462) group and the *Mllt11^flox/+^;Cre* (p=0.0302) group **(Fig. 5e).** We then visualized freezing of the three female genotypes specifically within each portion of the three-chamber task and found that, while the two-way ANOVA revealed there were no significant interactions of genotype and session on time spent freezing between the female genotypes (F(6, 93)=1.475; p=0.1953); there was a significant effect of session on time spent freezing (F(3,93)=3.095; p=0.0307). Multiple comparisons revealed that within the approach portion of the task, the cKO females spent significantly more time freezing than the *Mllt11^flox/+^* (p=0.0272) *and Mllt11^flox/+^;Cre* (p=0.0020) genotypes **(Fig. 5d)**. There were no significant differences between the average number of freezing bouts (p=0.0831; **Fig. 5g**), or the average bout length (p=0.5899; **Fig. 5h**) between the WT/Cre males. There were some significant differences when examining freezing at each portion of the test. There was a significant interaction effect of genotype and test portion (F(3, 36)=3.734; p=0.0196) and a significant effect of test portion (F(3, 36)=11.02; p<0.0001). Multiple comparisons revealed that the Cre animals spent significantly less time freezing than the WT animals in the Approach (p=0.0021), and Novelty (p=0.0475) portion of the three-chamber task **(Fig. 5i**). However, similar differences were not seen in our *Mllt11* heterozygous or cKO males **(Fig. 5f)**. Additionally, there was not a significant difference average number of bouts (p=0.4513; **Fig. 5j**) or the average length of freezing bout (p=0.3106: **Fig. 5k**) among the female WT/Cre control genotypes and no significant interaction effect of genotype and portion of the test on freezing time (F(3,39)=0.7293; p=0.5407; **Fig. 5l**) indicating that the effect on freezing we saw in our *Mllt11* cKO females was not due to the presence of Cre alone.

**Figure 5:**
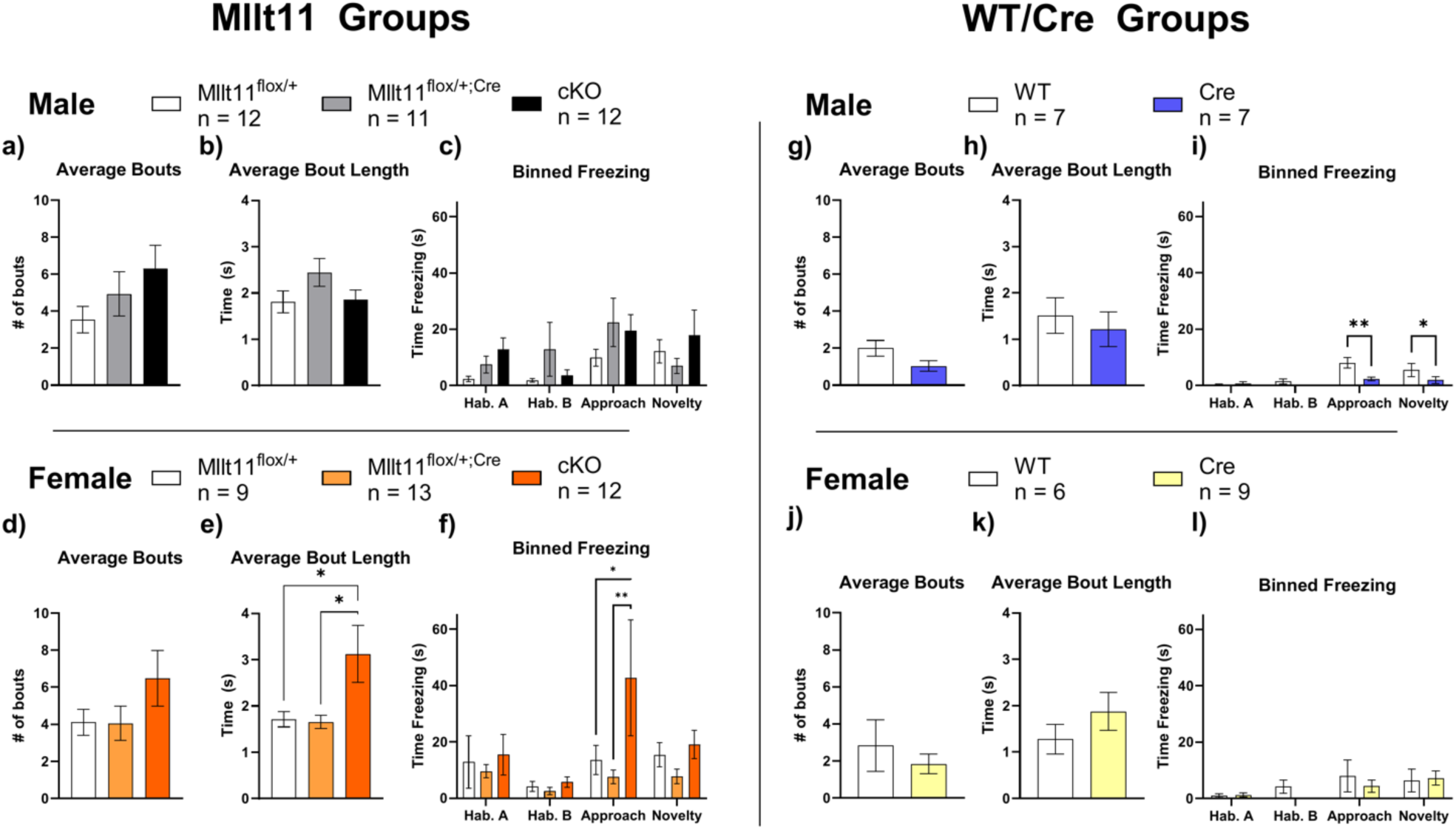
Three-Chamber Freezing. Graphical representation of the time spent freezing during the various stages of the three-chamber task. The average number of bouts across the entire three-chamber task was recorded and compared between the **a)** male *Mllt11^flox/+^, Mllt11^flox/+^;Cre*, and cKO genotypes, **d)** female *Mllt11^flox/+^, Mllt11^flox/+^;Cre*, and cKO genotypes, **g)** male WT and Cre genotypes, **j)** female WT and Cre genotypes. The average bout length across the entire three-chamber task was also recorded and compared between the **b)** male *Mllt11^flox/+^, Mllt11^flox/+^;Cre*, and cKO genotypes, **e)** female *Mllt11^flox/+^, Mllt11^flox/+^;Cre*, and cKO genotypes, **h)** male WT and Cre genotypes, **k)** female WT and Cre genotypes. To determine identify differential freezing behaviors across stages of the three-chamber task, the freezing in each stage of the test was presented independently. These sections were as follows: Habituation A: the time during which the mouse freely explored the center chamber while the doors to either side were closed; Habituation B: the time during which the mouse freely explored all three chambers; Approach: when the mouse could freely explore all three chambers, one of which now contained the ‘Stranger’; Novelty: when the mouse could freely explore all three chambers, one side of which contained the previous ‘Stranger’ mouse now referred to as the ‘Familiar’ mouse, and the other side of the chamber now contained the ‘Novel’ mouse. The time spent freezing was recorded and compared between the **c)** male *Mllt11^flox/+^, Mllt11^flox/+^;Cre*, and cKO genotypes, **f)** female *Mllt11^flox/+^, Mllt11^flox/+^;Cre*, and cKO genotypes, **i)** male WT and Cre genotypes, **l)** female WT and Cre genotypes. (For all graphs ns= not significant * = p<0.05; ** = p<0.01; *** = p<0.001; **** = p<0.0001).

Given the differences in freezing, we then investigated the exploratory locomotion of the mice via entry counts into any portion of the chamber (**Fig. 6)**. We did not find any significant differences between genotypes in the total number of entries made during any portion of the task for the males (F(4,64)= 1.721; p=0.1563) and all three male genotypes were more exploratory during the social approach (*Mllt11^flox/+^* p<0.0001; *Mllt11^flox/+^;Cre* p=0.0007 cKO p<0.0001) and novelty (*Mllt11^flox/+^* p<0.0001; *Mllt11^flox/+^;Cre* p<0.0001; cKO p<0.0001) portion of the task than during the habituation (**Fig. 6a**; significancy indicated by ###). Similarly, we did not find any significant differences between the three female genotypes (F(4, 62)= 0.8274; p=0.5128), and all three female genotypes were more exploratory during the social approach (*Mllt11^flox/+^* p=0.0002; *Mllt11^flox/+^;Cre* <0.0001; cKO p<0.0001) and novelty (*Mllt11^flox/+^*p=0.0002; *Mllt11^flox/+^;Cre* p<0.0001; cKO p<0.0001) portion of the task than during the habituation (**Fig. 6b**; significancy indicated by ###). The same was found to be true of the WT/Cre males (F(2,24)=2.852; p=0.0774; Approach: WT p<0.0001, Cre p<0.0001; Novelty: WT p=0.0059, Cre p<0.0001; **Fig. 6c**; significancy indicated by ##). For the Cre females, in addition to the increased exploratory behavior during approach (WT p=0.0002; Cre p=0.0003; **Fig 6d**; significancy indicated by ##) and novelty (WT p=0.0205; Cre p=0.0029; **Fig 6d**; significancy indicated by ##) for both control genotypes (**Fig. 6d**), there was a significant interaction effect of genotype and test portion on number of entries (F(2,26)=4.747; p=0.0174) with multiple comparisons revealing that the WT females were more exploratory than the Cre females during the approach portion (p=0.0229; **Fig 6d**; significancy indicated by *). We then binned the entries data for the approach portion of the task for the WT/Cre females to understand this difference. The WT females made significantly more entries than the Cre females into the empty chamber (p=0.0142; **Fig. 6e**; significancy indicated by *) and the center chamber (p=0.0223; **Fig. 6e**; significancy indicated by *), but there were no differences in the number of entries to the stranger chamber (p=0.0780; **Fig. 6e**). This correlates with our previous results showing the WT females spending more time in the empty side than with the stranger and indicates that as their number of entries into the stranger side did not differ between genotypes that the differences in time spent in the stranger chamber are due to time alone and not lack of exploration into that chamber by the WT animals. In addition, these results indicate that the differences in freezing we found, particularly in *Mllt11* cKO females, did not impact the overall exploratory behaviors of the animals; finally, the differences in the WT/Cre animals were also not replicated in the *Mllt11* heterozygous and homozygous genotypes. Thus, we can conclude that the differences in preference and investigation times were not due to any generalized lack of ambulatory exploration or any effect of Cre on exploratory behaviors.

**Figure 6:**
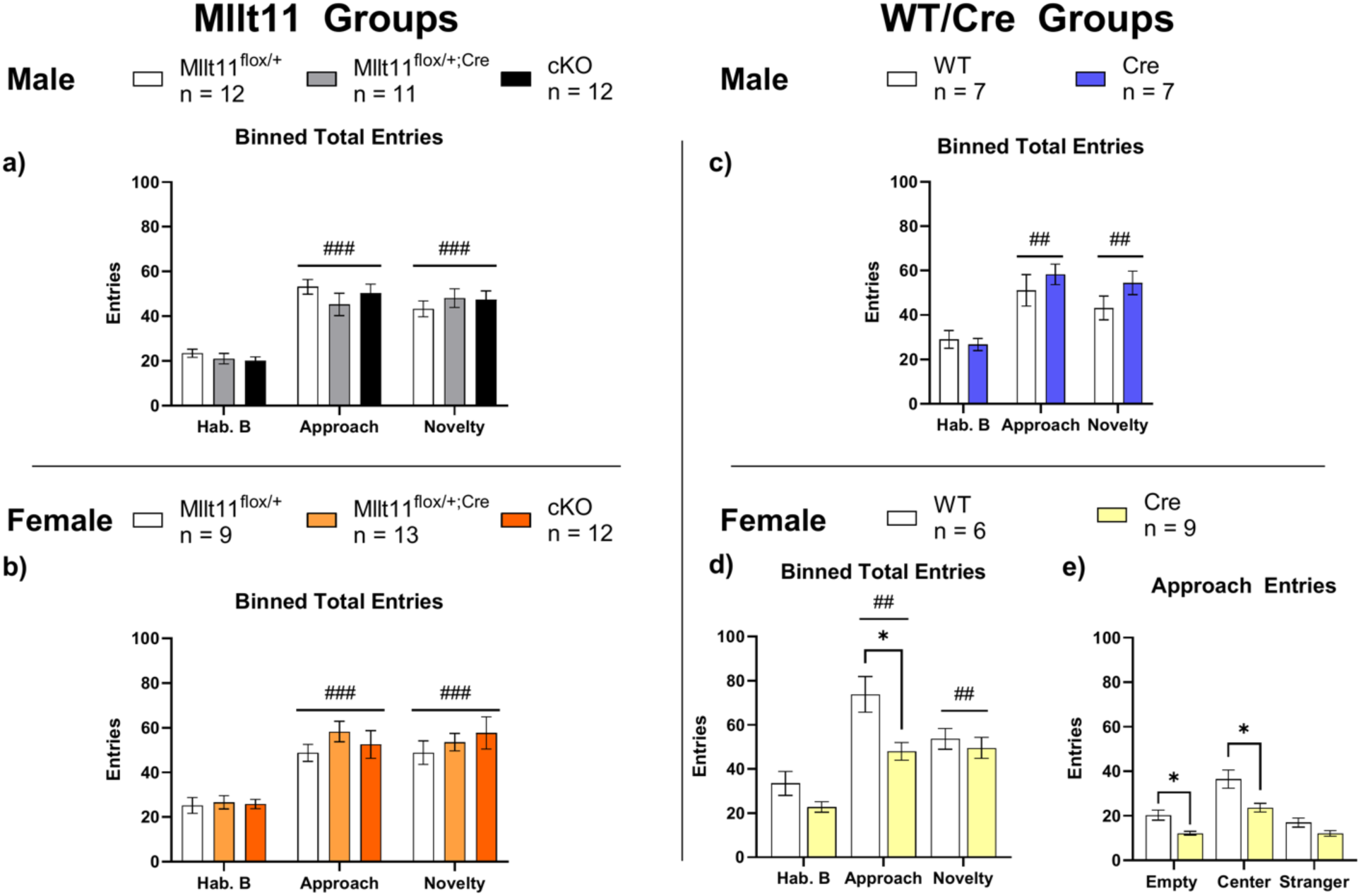
Three-Chamber Entries. Representation of the number of entries during the portions of the three-chamber task in which the doors were open. **a)** Total entries compared between the male *Mllt11^flox/+^, Mllt11^flox/+^;Cre*, and cKO genotypes during Habituation B, Social Approach, and Social Novelty. (### - indicates that all three genotypes are significantly different from their values in Habituation B) **b)** Total entries compared between the female *Mllt11^flox/+^, Mllt11^flox/+^;Cre*, and cKO genotypes during Habituation B, Social Approach, and Social Novelty. (### - indicates that all three genotypes are significantly different from their values in Habituation B) **c)** Total entries compared between the male WT and Cre genotypes during Habituation B, Social Approach, and Social Novelty. (## - indicates that all two genotypes are significantly different from their values in Habituation B) **d)** Total entries compared between the female WT and Cre genotypes during Habituation B, Social Approach, and Social Novelty. (## - indicates that all two genotypes are significantly different from their values in Habituation B) **e)** Binned entries into the empty, center, or stranger side during the Social Approach portion of the task for the WT and Cre females. (### - indicates that all three genotypes are significantly different from their values in Habituation B; ## - indicates that all two genotypes are significantly different from their values in Habituation B; For all graphs ns= not significant * = p<0.05; ** = p<0.01; *** = p<0.001; **** = p<0.0001).

### Female cKO animals display reduced nestlet shredding

On P51, we performed both a marble burying task and a nestlet shredding task. One-way ANOVAs found no significant differences in number of marbles buried between the male experimental genotypes (*Mllt11^flox/+^*, *Mllt11^flox/+^;Cre*, and cKO; F(2,31)=1.178; p=0.3213; **Fig. 7a**). After the animals returned to their home cage in the animal colony for one hour, we performed a nestlet shredding task. The males did not display any difference in percent of nestlet shredded (F(2,31)=0.3958; p=0.6765; **Fig. 7b**). Similarly, we found no significant differences in number of marbles buried between the female experimental genotypes (*Mllt11^flox/+^*, *Mllt11^flox/+^;Cre*, and cKO; F(2,29)=0.7560; p=0.4786; **Fig. 7c**). For the nestlet portion however, one-way ANOVA revealed a significant effect in the females (F(2,31)=5.279; p=0.0106) and Tukey’s multiple comparisons revealed that the female cKO animals shredded significantly less nestlet than the *Mllt11^flox/+^*females (p=0.0089; **Fig. 7d**). We then confirmed that the effects we saw were not due to Cre expression alone via analysis of the WT/Cre genotypes. Similarly, we found no significant differences in the number of marbles buried by the male (p=0.9364; **Fig. 7e**) or the amount of nestlet shredded (p=0.2127; **Fig. 7f**). Between the WT/Cre female control genotypes, we found no significant difference in number of marbles buried (p=0.7041; **Fig. 7g**) and we confirmed that the reduction in nestlet shredding was not due to Cre expression as the female WT and Cre animals did not display any significant differences in the percentage of nestlet shredded (p=0.2435; **Fig. 7h**).

**Figure 7:**
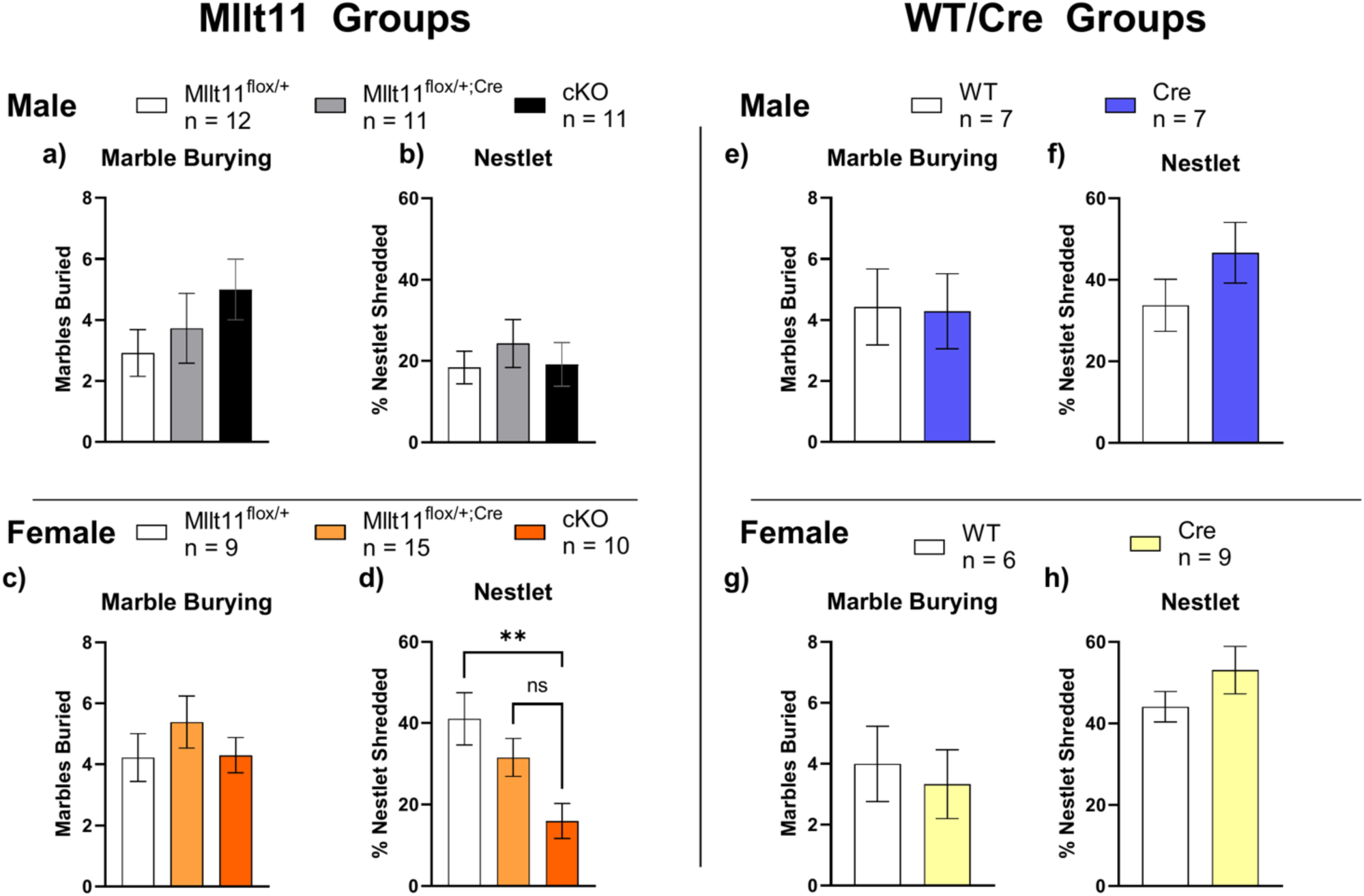
Marbles and Nestlet Shredding. Graphical representation of the tests performed on P55 which included the buried marble test followed by the nestlet shredding task. The number of marbles buried following 30 minutes in the rat cage was recorded and compared between the **a)** male *Mllt11^flox/+^, Mllt11^flox/+^;Cre*, and cKO genotypes, **c)** female *Mllt11^flox/+^, Mllt11^flox/+^;Cre*, and cKO genotypes, **e)** male WT and Cre genotypes, **g)** female WT and Cre genotypes. The percentage of the nestlet square was recorded following 30 minutes and compared between the **b)** male *Mllt11^flox/+^, Mllt11^flox/+^;Cre*, and cKO genotypes, **d)** female *Mllt11^flox/+^, Mllt11^flox/+^;Cre,* and cKO genotypes, **f)** male WT and Cre genotypes, **h)** female WT and Cre genotypes. (For all graphs ns= not significant * = p<0.05; ** = p<0.01; *** = p<0.001; **** = p<0.0001).

### Mllt11 loss did not result in presentation of repetitive behaviors

As repetitive and stereotypic behaviors are a core phenotype of ASD, we desired to examine our mice for presentation of any repetitive behaviors. We measured the amount of time spent grooming, digging, and rearing within the P30 olfaction habituation/dishabituation task. There were no significant differences for the *Mllt11^flox/+^*, *Mllt11^flox/+^;Cre*, and cKO males in grooming (F(2,31)=0.4023; p=0.6722; **Fig. 8a**), digging (F(2,31)=0.4928; p=0.6156; **Fig. 8b**); or rearing (F(2,31)=1.648; p=0.2089; **Fig. 8c**). Likewise, within the *Mllt11^flox/+^*, *Mllt11^flox/+^;Cre*, and cKO females there were no significant differences in grooming (F(2,34)=1.892; p=0.1663; **Fig. 8d**), digging (F(2,34)=1.647; p=0.2076; **Fig. 8e**), or rearing (F(2,34)=0.4441; p=0.6451; **Fig. 8f**). While there was no difference in male WT/Cre grooming (p=0.3482; **Fig. 8g**), or digging (p=0.2980; **Fig 8h**), the male Cre controls did rear significantly more than the WTs (p=0.0302; **Fig. 8i**). Among the female WT/Cre control genotypes, there were no significant differences between genotypes for grooming (p=0.3084; **Fig. 8j**) but there was a significant difference in digging as the Cre females spending significantly more time digging than the WT females (p=0.0131; **Fig. 8k**). Finally, there was not a significant difference between the Cre/WT females and the number of rears (p=0.5781; **Fig. 8l**).

**Figure 8:**
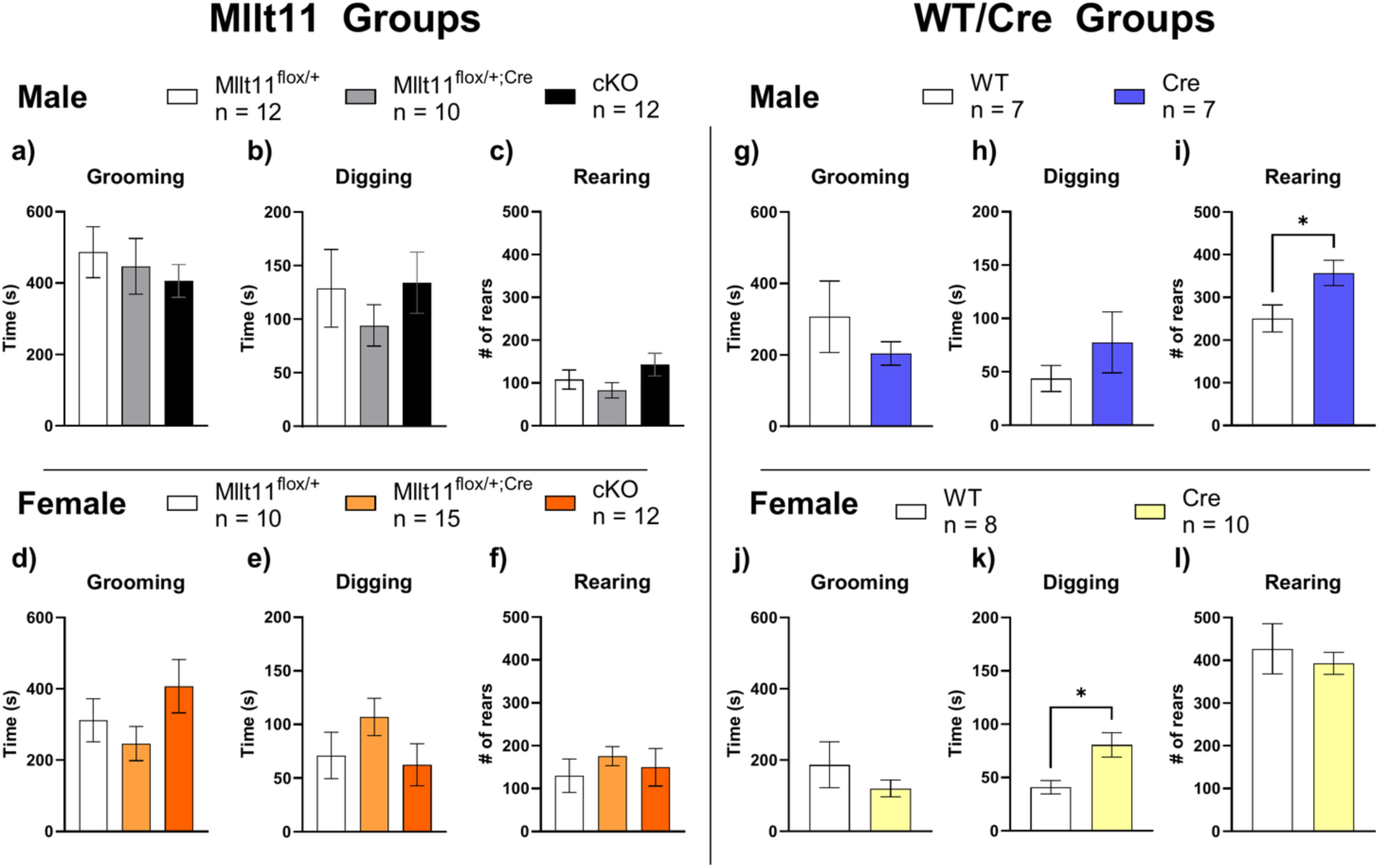
Repetitive Behaviors. Graphical representation of the repetitive behaviors scored via video that occurred during the Olfaction Habituation/Dishabituation task performed on P30. These included grooming, digging, and rearing. **a-c)** male *Mllt11^flox/+^, Mllt11^flox/+^;Cre*, and cKO genotypes, **d-f)** female *Mllt11^flox/+^, Mllt11^flox/+^;Cre*, and cKO genotypes, **g-i)** male WT and Cre genotypes, **j-l)** female WT and Cre genotypes. (For all graphs ns= not significant * = p<0.05; ** = p<0.01; *** = p<0.001; **** = p<0.0001).

### Histology: Reduced Corpus Callosum and Cortical Thickness

In humans, the best-replicated structural brain phenotype found in individuals with autism is reduced corpus callosum size^34^. Reduced corpus callosum projections and cortical thickness have been seen in this mouse model at embryonic timepoints but have not been examined postnatally^8^. We thus examined the corpus callosum width and cortical thickness in a cohort of animals who did not undergo behavioral analysis. Animals were aged to P30 to coincide with the onset of our observed behavioral phenomena and then transcardially perfused. Brain tissue was collected, immunohistochemically stained for neurofilament (NF) a neuronal-specific intermediate filament found in axons, and images were taken (Representative images: **Fig 9a-d**). For each individual, three measurements of the corpus callosum were taken and the average of those three measurements was analyzed (**Fig. 9aii**: red dashed line indicates were each measurement was taken per image (3 images per individual)). One-way ANOVAs found significant differences in corpus callosum thickness between the male experimental genotypes (*Mllt11^flox/+^*, *Mllt11^flox/+^;Cre*, and cKO) (F(2,12)=20.66; p=0.0001); **Fig. 9e**). Tukey’s multiple comparisons revealed that both the cKO mice and the *Mllt11^flox/+^;Cre* genotypes had significantly thinner corpus callosum than the *Mllt11^flox/+^* mice (p=0.0001; p=0.0032), but no difference in thickness was found between the *Mllt11^flox/+^;Cre* and cKO genotypes (p=0.1328; **Fig. 9e).** Cortical thickness measurements were acquired from each individual with three images being acquired per individual, then three measurements per image were acquired of the cortical thickness (**Fig 9aV**: red dashed lines indicate an example of three measurements taken per image (3 images per individual). The average of those 9 measurements was then analyzed. Examination of cortical thickness with One-way ANOVA found significant differences in cortical thickness between the male experimental genotypes (*Mllt11^flox/+^*, *Mllt11^flox/+^;Cre*, and cKO) (F(2,12)=5.630; p=0.0189); **Fig. 9f**). Tukey’s multiple comparisons revealed that the cKO mice had significantly thinner cortex measurements than the *Mllt11^flox/+^* mice (p=0.0149; **Fig. 9f**).

**Figure 9:**
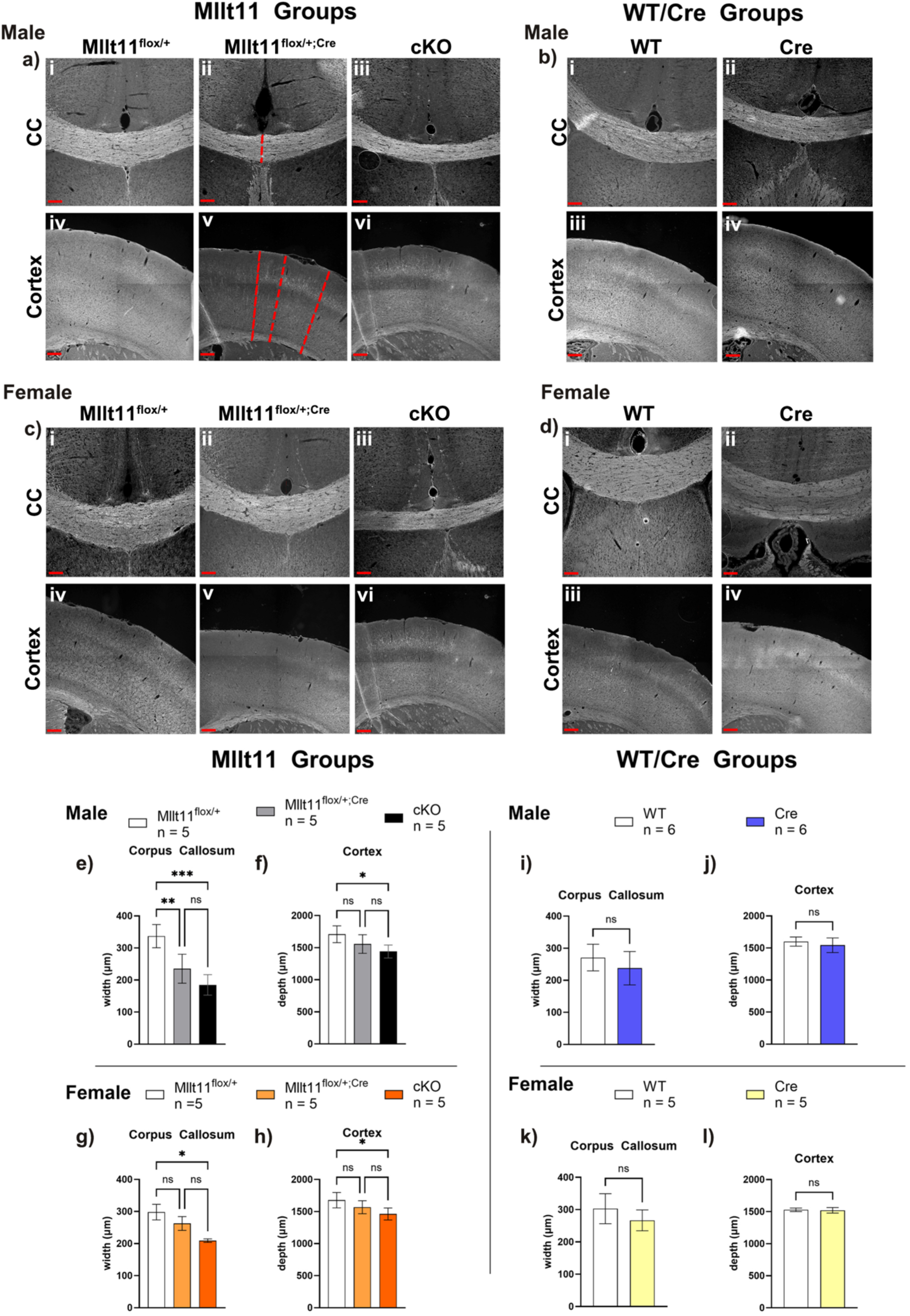
Cortical and Corpus Callosum Thickness. Representative images and graphical representation of the cortical and corpus callosum measurements **a)** Representative images of the corpus callosum from male **(i)** *Mllt11^flox/+^,* **(ii)** *Mllt11^flox/+^;Cre*, and **(iii)** cKO genotypes. White: Neurofilament; Scale bar = 100µm. Dashed lines in (**ii)** and **(v)** demonstrate the measurements taken for the corpus callosum width and cortex depth. For the corpus callosum one measurement was taken per image (3 images per animal); for the cortex, three measurements were taken per image (3 images per animal). Representative image of the cortex from male **(iv)** *Mllt11^flox/+^,* **(v)** *Mllt11^flox/+^;Cre*, and **(vi)** cKO genotypes. White: Neurofilament; Scale bar = 200µm. **b)** Representative images of the corpus callosum from female **(i)** *Mllt11^flox/+^,* **(ii)** *Mllt11^flox/+^;Cre*, and **(iii)** cKO genotypes. White: Neurofilament; Scale bar = 100µm. Representative image of the cortex from female **(iv)** *Mllt11^flox/+^,* **(v)** *Mllt11^flox/+^;Cre*, and **(vi)** cKO genotypes. White: Neurofilament; Scale bar = 200µm. **c)** Representative images of the corpus callosum from male **(i)** WT, **(ii)** Cre White: Neurofilament; Scale bar = 100µm. Representative image of the cortex from male **(iii)** WT **(iv)** Cre. White: Neurofilament; Scale bar = 200µm. **d)** Representative images of the corpus callosum from female **(i)** WT, **(ii)** Cre. White: Neurofilament; Scale bar = 100µm. Representative image of the cortex from male **(iii)** WT **(iv)** Cre White: Neurofilament; Scale bar = 200µm. **e-l**) Graphical representation of measurements acquired from brain tissue of mice at post-natal day 30. Corpus callosum thickness was compared between **e)** male *Mllt11^flox/+^, Mllt11^flox/+^;Cre*, and cKO genotypes, **g)** female *Mllt11^flox/+^, Mllt11^flox/+^;Cre*, and cKO genotypes, **i)** male WT and Cre genotypes, and **k)** female WT and Cre genotypes. Cortical thickness was compared between **f)** male *Mllt11^flox/+^, Mllt11^flox/+^;Cre*, and cKO genotypes, **h)** female *Mllt11^flox/+^, Mllt11^flox/+^;Cre*, and cKO genotypes, **j)** male WT and Cre genotypes, and **l)** female WT and Cre genotypes. (For all graphs ns= not significant * = p<0.05; ** = p<0.01; *** = p<0.001; **** = p<0.0001).

The same examination between the female experimental genotypes (*Mllt11^flox/+^*, *Mllt11^flox/+^;Cre*, and cKO) found significant differences in corpus callosum thickness (F(2,12)=5.526; p=0.0199; **Fig. 9g**) with Tukey’s multiple comparisons revealing that the cKO mice had significantly thinner corpus callosum than the *Mllt11^flox/+^* mice (p=0.0161). Similarly to the males, examination of cortical thickness with One-way ANOVA in the females found significant differences (F(2,12)=5.203; p=0.0236; **Fig. 9h**). Tukey’s multiple comparisons showed that the cKO mice had significantly thinner cortex measurements that the *Mllt11^flox/+^* mice (p=0.0185; **Fig. 9h**).

To determine whether these results could be due to the presence of cre alone, we examined the corpus collosum and cortical thickness between the control genotypes as well. No significant differences were found between the Cre/WT males in regard to corpus callosum thickness (p=0.2509; **Fig. 9i**) or cortical thickness (p=0.3226; **Fig. 9j**). Similarly, no differences were found between the Cre/WT females in regard to corpus callosum thickness (p=0.1919; **Fig. 9k**) or cortical thickness (p=0.8793; **Fig. 9l**).

## DISCUSSION

The current study revealed that *Mllt11* loss in mice resulted in sex-dependent behavioral alterations. Specifically, we found the female *Mllt11* cKO mice had reduced interest in social odors, however, when in the presence of social cues elicited by the physical presence of another mouse in the three-chamber task, the cKO females displayed the expected preference for the social side of the chamber as well as investigation of the stranger mouse. Interestingly, they then did not show a preference for social novelty in the final portion of the three-chamber task. The same effects were not seen in the Cre control female cohort, indicating these effects were specifically due to the loss of *Mllt11*. *Mllt11* cKO males, on the other hand, showed no disruption in social interest during the olfaction test, as well as normal social approach. However, only those lacking one or both copies of *Mllt11* investigated the novel mouse significantly more than the familiar mouse. Future studies will need to perform novel object recognition tasks to investigate the perception and preference for novelty in both the male and the female mice.

While the females did not show any significant differences in the marble burying task or in any measures of behaviors such as grooming, digging, or rearing, the female cKOs did shred significantly less nestlet. In the literature, excessive shredding is an indicator of repetitive behavior; however, reduced nestlet shredding has been viewed as restricted interest in novel objects^35^. In combination with the lack of preference for social novelty, we believe this is best interpreted as restricted interest in novel objects or decreased responsivity to novelty^35^. This would align with a symptom seen in children with ASD as some have been found to not show attentional preferences for novel stimuli under certain conditions^36^. Novel object recognition tasks should be performed in future studies to identify whether this interpretation is correct in our *Mllt11* cKO animals.

Finally, the higher total amount of freezing exhibited by the female cKO animals during the three-chamber task with no increase in marble burying may be indicative of social context-specific freezing. Specifically, we saw that the female knockouts spent significantly more time freezing when the first stranger was added to the chamber. These differences were driven not by the number of freezing bouts, but by the length of time spent freezing. An increase in marble burying as well would be expected if there were a broad underlying higher baseline anxiety – thus we hypothesize that the increased freezing seen in the cKO female animals may be increased anxiety specifically in the social context. This would correlate with human findings as higher social anxiety is seen in human females as opposed to human males with ASD^37^. However, specific social and non-social anxiety-related tests will need to be performed to determine the validity of this hypothesis.

### Mechanisms and Sex Differences

Sex differences in NDD presentation are well documented^38^. For example, in ASD, females experience more severe internal symptoms such as anxiety and depression and, while both males and females have disrupted social and communication skills. It is also suggested that females ‘camouflage’ their social deficits more than their male counterparts^39^. For this reason, we find it particularly intriguing that the female cKOs displayed reduced interest in social odors but no differences in social preference or investigation time in a more social condition like the three-chamber social approach task. We hypothesize that the strength of the available social cues could contribute to these differences. The saliency of a swab from a dirty cage is expected to be lower than that of the physical presence of another mouse. As such, if the social disturbances are mild, one might expect the presence of more salient social cues to override any deficits. Indeed, in humans with ASD, it has been shown that the saliency of social cues is perceptible and affects attention despite an overall deficit in social attention^40^. Future studies should investigate the ultrasonic vocalizations of these mice to better understand what degree of social deficits may exist.

Regarding mechanism, these current results demonstrate that the corpus callosum thickness and cortical thickness were reduced in the *Mllt11^flox/+^;Cre* and cKO males as well as the cKO females at P30; however, how this same effect then results in sex-specific behavioral differences is unclear. The impact of Mllt11 on maintenance of UL2/3-specific markers may provide a possible mechanism through which it may play a role in modulation of behavior. Previous work by our lab found that the loss of *Mllt11* resulted in decreased expression of Satb2 as well as CDP/Cux1 in the superficial somatosensory cortex^8^. First, CDP/Cux1 has been shown to play a role in regulating both neurite outgrowth and overall neuronal morphology, and its loss gives rise to neurons with shorter, less arborescent neurites^41,42^. Neuronal morphology altered in this manner has been established as a hallmark feature of both human and mouse ASD or ASD-like pathology^43,44^. Indeed, we previously reported that *Mllt11* cKO results in reduced callosal crossing fibers and less complex arborization patterns at embryonic timepoints^8^. There is some evidence in the literature from animal models that indicates sex-dependent disruptions of Cux1 expression following prenatal exposure to bisphenol A (BPA)^45^, an implicated risk factor for NDDs^46^. Second, Satb2 mutation in humans gives rise to SATB2 syndrome, an NDD characterized by intellectual disability, language and communication deficits, some ASD-like behaviors such as repetitive or restrictive movements and interests, as well as hyperactivity and aggression^47^. However, the effects of *Mllt11* cKO were mainly seen in our female mice and to date no sex-differences have been reported in Satb2 syndrome^48,49^. Future studies should examine whether the expression of Satb2 and CDP/Cux1 are reduced following *Mllt11* loss in a sex-dependent manner, as our previous work did not consider sex as a variable.

Another hypothesis is that the sex-specific effects may be due to two potential Mllt11 interactions, including Wnt signaling and non-muscle myosin NMIIA/B interactions. To begin, Mllt11 is known to activate Wnt signaling and is specifically implicated in the activation of the β-catenin-mediated canonical Wnt pathway^50^. It is hypothesized that the level of Wnt/β-catenin signaling could contribute to the phenotypical heterogeneity of ASD^51^. Research on the activation of the Wnt pathway in other NDDs supports this hypothesis. One such example in the case of SZ is expression of DEK, a chromatin-remodeling phosphoprotein which, like Mllt11, is an oncogene^52^ and activator of the Wnt pathway^53^. O’Donovan, Franco-Villanueva, Ghisays, Caldwell, Haroutunian, Privette Vinnedge, McCullumsmith and Solomon ^53^ found that DEK protein depletion down-regulated the Wnt pathway and was a marker of cognitive impairment. However, these effects were sex-specific such that lower DEK, and hypothetically less down-regulation of the Wnt pathway, in females was associated with more severe cognitive impairment, whereas in males, higher levels of DEK were associated with severity of cognitive impairment. As Mllt11 is known to activate Wnt signaling^50^, we hypothesize that loss of *Mllt11* may alter Wnt/β-catenin signaling and differentially alter behavior thus contributing to the phenotypical heterogeneity of ASD^51^.

Previous work from our lab has shown that Mllt11 interacts with two non-muscle myosin II (NMII) paralogs, NMIIA and NMIIB^10^. Furthermore, we found that *Mllt11* cKO resulted in increased expression of NMIIB ^10^. Intriguingly, NMIIB overexpression is known to inhibit the Wnt/β-catenin pathway^54^ while NMIIA activates Wnt/β-catenin signaling by interacting with β-catenin and controlling β-catenin transcriptional activity^55^. In mice, NMIIB was found to be differentially expressed in the brains of male and female mice and is a known ASD risk gene^56^. With this being the case, we hypothesize that the combined differential expression of NMIIA/B and its differential contribution to the Wnt/β-catenin pathway could contribute to the sex-specific disruptions. Future experiments should determine if *Mllt11* loss differentially regulates the Wnt/β-catenin pathway, whether the increased expression of NMIIB is found in both males and females, and whether these alterations are present consistently across the lifespan of the mice.

Finally, there is a general belief that the estrous cycle results in high levels of variance in female rodents. While it is well established that the cyclical fluctuations in hormones do alter sexual behaviors of female rodents^57,58^, recent work suggests that behavioral variations in females is generally not different than the variations in males^59–63^. Due to the additional animals required to achieve sufficient n’s within each cycle stage, as well as the added stress that might impact behaviour through cycle monitoring protocols, we decided to not monitor the estrous cycles of our female animals.

### *Cux2^iresCre^* Contributions to NDD Behavioral Phenotypes

To our knowledge, we are the first to report behavioral effects due to *Cux2^iresCre^* expression. Our Cre control genotypes displayed several significant differences that were unexpected. Our male Cre mice did not spend a significantly greater amount of time in the chamber or investigating the stranger mouse as opposed to the empty pencil cup. As we saw a similar lack of significance in our *Mllt11^flox/+^;Cre*, and cKO males, this suggests that these particular effects were not solely due to the loss of *Mllt11*. The female Cre mice displayed a preference for the stranger mouse, both in time spent in the chamber and investigation time. However, the WT females displayed a preference for the empty chamber and did not investigate the stranger more than the empty pencil cup. These results are perplexing, and we included an additional measurement of freezing to see if this may have been the source of the effect. However, we did not see an increase in freezing behavior of the WT females, suggesting this lack of preference was not due to freezing. Further testing will be required to parse apart the source of this effect. However, since we did not see a similar effect in our female *Mllt11^flox/+^*, *Mllt11^flox/+^;Cre*, and cKO genotypes, we believe this is a difference that does not impact the results of our study. We also found that the female Cre mice spent significantly more time digging; however, this result was not reflected in our female *Mllt11^flox/+^*, *Mllt11^flox/+^;Cre*, and cKO genotypes. Cre lines are frequently utilized with the assumption that the expression of the Cre gene results in either minimal or no phenotypic behavioral changes. Unfortunately, this has been shown to not be the case by many groups and within various Cre mice including ChAT-Cre lines^64,65^, DAT-Cre ^66,67^, Nestin-Cre^68^. Others have also found sex-specific effects in other Cre mice^69^. Specifically, Baghdadi, Mesaros, Purrio and Partridge ^69^ reported that when examining male and female Syn1Cre mice, they found a male-specific increase in anxiety-like behaviors and reduced body weight^69^. Interestingly, in the liver of mice and rats, Cux2 is expressed at significantly higher levels in females than in males^70^ but whether sex-differences in Cux2 expression are present in the brain is unknown. For future studies, we believe it is important to perform more extensive behavioral testing of the *Cux2^iresCre^* mice to determine if replication of these differences is possible and to investigate the contribution of Cre expression to these differences.

Another consideration when interpreting the behavioral phenotypes of mutant mouse models of NDDs is the background of the strain in which the mutation is maintained. Our line was maintained in a mixed C57Bl/6J/FVB/NJ background. C57Bl/6 mice have been shown, in some cases, to display various autism-like behaviors under wild type conditions. This phenomenon is likely due to the level of inbreeding required in maintaining the genetic purity characteristic of this strain^71^. Alternatively, genetic outbreeding and the genetic variability enhanced when crossing strains may also be a source of variability in behaviors. As the mice used in the experiments reported herein were derived from outbred C57Bl/6/FVB hybrid crosses, this requires careful consideration when parsing the impact of *Mllt11* loss from other potential sources of variation. To try to mitigate the effects of this variability, littermates of all genotypes were used as behavioral controls. For this reason, we believe that the littermate *Mllt11^flox/+^* mice are a better control than the WT mice which were not littermates.

## CONCLUSION

In conclusion, we found that our female *Mllt11* cKO mice display more ASD-like behavioral deficits than our male *Mllt11* cKO mice, including reduced interest in social odors and reduced nestlet shredding. We also confirmed that the reduction in thickness of both the cortex and corpus callosum seen at embryonic timepoints in this mouse model persists in postnatal mice. As 80% of those diagnosed with autism are male children, pre-clinical investigations typically focus specifically on male mice. This focus can have, as Murta, Seiffe and Depino ^72^ have argued, “detrimental consequences on our understanding of ASD etiology and pathophysiology.” We therefore believe that the presentation of a mouse model with ASD-like behavioral deficits in the females but not the males is intriguing and necessary to utilize in future investigations. Future studies are needed to identify the depth of the communication deficits as well as the perception of novelty and identification of the molecular contributions to the sex-differences we observed.

## Author contributions

EW and DST designed and conducted experiments, scored and analyzed behavior, and co-wrote the manuscript; PG and JG scored behavior; AI edited the manuscript, secured funding, and supervised project.

## Acknowledgements

Funding provided by the Canadian Institutes of Health Research (PJT-388914), and the Killam Foundation and American Association for Anatomy for post-doctoral fellowship support to EW. We thank Sarah Whitehead for assistance with animal husbandry.

